# Stress granule sequestration of OXPHOS gene transcripts exacerbates glycolytic restriction

**DOI:** 10.1101/2025.09.02.673658

**Authors:** Wanling Zheng, Ruoqing Xu, Maoguang Xue, Xiaoyu Liu, Yinglong Gao, Min-Xin Guan, Jun Ma, Feng He

## Abstract

Stress granules (SGs) are dynamic organelles formed under cellular stress and they are generally regarded as protective entities. Meanwhile, their role in pathogenesis is becoming increasingly recognized, but the underlying mechanisms remain elusive due to the diverse nature of both stress types and biological contexts. Here we investigate SG dynamics and temporal changes in bulk and SG-associated transcriptomes under different regimens that inhibit glycolysis. We subject cells to either single assaults of glucose depletion (GD) or 2-deoxy-D-glucose addition (2DG) or a combined treatment (GD+2DG). We find that SGs formed under these conditions exhibit distinct properties, including eIF2α phosphorylation dependency, mRNA composition, and capacity to disassembly. Our results show that SGs induced by GD+2DG uniquely trap oxidative phosphorylation (OXPHOS) gene transcripts, leading to mitochondrial dysfunction. We provide evidence suggesting that the persistency of SGs formed under GD+2DG treatment is interwoven with mitochondrial dysfunction resulting in heightened apoptosis, effects that can also be recreated under single assaults when combined with mitochondrial inhibition. Our findings suggest that SG formation induced by inhibiting a single metabolic pathway can widen its impact in intensifying cellular metabolic stress under specific conditions, providing mechanistic insights into the paradoxical dual nature of SGs in stress response and pathology.

## Introduction

Eukaryotic cells respond to stress by compartmentalizing translationally aberrant ribonucleoproteins into stress granules (SGs), which are dynamically assembled and disassembled to regulate mRNA and protein availability (Van Leeuwen et al. 2019; Campos-Melo et al. 2021). This process enables cells to conserve energy, survive stress, and recover efficiently. However, dysregulation in SG assembly or disassembly can render cells vulnerable to stress, contributing to the pathogenesis of various diseases, including amyotrophic lateral sclerosis (ALS) and frontotemporal dementia (FTD) (Zhang et al. 2019; Parameswaran et al. 2023; Buchan et al. 2013; Cui et al. 2023; Wolozin et al. 2019; Cui et al. 2024). Understanding the molecular mechanisms governing SG dynamics is therefore critical to elucidation of the underlying causes of these disorders.

SG formation as a cellular process is mediated by diverse signaling pathways molecularly, depending on the precise nature and context of the stress. For instance, under arsenite-induced oxidative stress or heat shock, SG assembly is primarily triggered by the phosphorylation of the translation initiation factor eIF2α (Frydrýšková et al. 2020; Aulas et al. 2017; Szczerba et al. 2023). Beyond this canonical pathway, alternative mechanisms, such as the activation of 4EBP1 and disruption of the eIF4F complex, have also been implicated in SG formation under conditions such as selenite exposure, hydrogen peroxide treatment, or complete glycolysis inhibition (Tauber et al. 2020; Emara et al. 2012; Wang et al. 2022; Fujimura et al. 2012). These findings highlight both versatility and specificity of SG responses to cellular stress.

The characteristics of SGs can also be influenced by the intensity and duration of the stress. Time-course analyses reveal that under arsenite exposure, the size and dynamics of the SG core, marked by the G3BP1 protein, remain stable over 2 hours, suggesting minimal changes in their biochemical state (Wheeler et al. 2016). In contrast, heat shock-induced SGs exhibit significant changes in their core size over time, accompanied by a proteomic transition as the stress prolongs (Mateju et al. 2017; Hu et al. 2023). Furthermore, the dynamics of SG dissolution upon alleviation of the stress or cellular adaptation to the stress may also rely on context-specific mechanisms, which can involve distinct signaling pathways, molecular chaperons, and macromolecule degradation systems (Jia et al. 2024; Hofmann et al. 2021). However, most of the existing studies have focused on acute stress-induced SGs, and a comprehensive knowledge is currently lacking with regard to the dynamics and composition of SGs formed under chronic stress (Hofmann et al. 2012; Cherkasov et al. 2013; Huang et al. 2020; Reineke et al. 2018, 2019; Youn et al. 2018, 2019; Khong et al. 2017; Namkoong et al. 2018; Somasekharan et al. 2020; Frydrýšková et al. 2020).

Nutrient starvation and mitochondrial inhibition have emerged as powerful paradigms for studying chronic stress-induced SGs (Reineke et al. 2018; Fu et al. 2016; Eiermann et al. 2022; Wang et al. 2022; Aguilera-Gomez et al. 2017; Amen et al. 2021; Sfakianos et al. 2018; Pernin et al. 2024). For example, when glucose, serum, glutamine and pyruvate were deprived all together, SGs assembled slowly, reached peak formation at 8 hours, persisted at 16 hours, and disassembled upon addition of glucose (Amen et al. 2021; Reineke et al. 2018). However, existing models are complicated by the fact that energy depletion—due to either nutrient withdrawal or mitochondrial dysfunction—not only represents a specific type of stress but also impairs the cell’s ability to assemble and disassemble SGs in response to other stressors (Jain et al. 2016; Pernin et al. 2024; Wang et al. 2022; Eum et al. 2020). Here we investigate mechanisms underlying SG dynamics under glycolytic inhibition, aimed at disentangling these complexities to provide insights into chronic stress responses.

In this study, we compare SG dynamics under different long-term glycolytic inhibition regimens. We show that while treatments of glucose depletion (GD) or 2- deoxy-D-glucose (2DG) addition induce SGs that dissipate as stress prolongs from 8 to 24 hours, the combined GD+2DG treatment results in a persistent presence of SGs and an increase in apoptotic cells over time. Time-resolved bulk RNA-seq reveals a unique upregulation of oxidative phosphorylation (OXPHOS) pathway activity under GD+2DG. However, this upregulation seems to be a futile response since transcripts of many OXPHOS genes are specifically sequestered within SGs, as evidenced by G3BP1-APEX2-enriched RNA-seq. We verify a reduced expression of OXPHOS proteins and mitochondrial dysfunction, properties that differentiate cells that are treated with GD+2DG from those under single treatments. We suggest that mitochondrial defects in these cells were responsible for SG persistence, and such SG persistency can be recapitulated in cells under single glycolytic inhibition with mitochondrial disruption. Our findings support a feedback model in which SG formation itself has a role in widening the impact of glycolytic restriction through limiting OXPHOS gene expression, creating a scenario of a deepened, perpetuating cellular stress.

## Results

### GD, 2DG and GD+2DG induce SGs with distinct formation kinetics

The aim of this study was to compare how cells may form stress granules (SGs) in response to different assaults that inhibit a common metabolic pathway, glycolysis. Here we investigated this question through subjecting 143B cells to either single or combined assaults. For single assaults, cells were treated with glucose depletion (GD) or with addition of 2-deoxy-D-glucose (2DG), a glucose analog that directly blocks glycolysis. For the double assault, 2DG was supplemented while glucose was withdrawn from the medium (GD+2DG). We examined the characteristics of SG core during their formation using G3BP1 as the marker (Figure 1A). ∼20% of the cells under the treatment of GD or 2DG became SG-positive (SG+) at the peak time of 8 h and 1 h, respectively, while 75% cells were SG+ under the treatment of GD+2DG at the peak time of 1 h (Figure 1B). The number of SGs per cell and the average size of SGs were comparable between the two single-assault groups but distinct from the GD+2DG group (Figure 1C-D). Based on these SG characteristics and those detailed in the following sections, we regard GD as chronic-mild stress, 2DG as acute-mild stress, and GD+2DG as acute-severe stress. As explained further below, we propose that the previously reported eSG (Wang et al. 2022) is a result of a further exacerbated stress state that goes beyond a mere glycolysis restriction.

**Figure 1.**
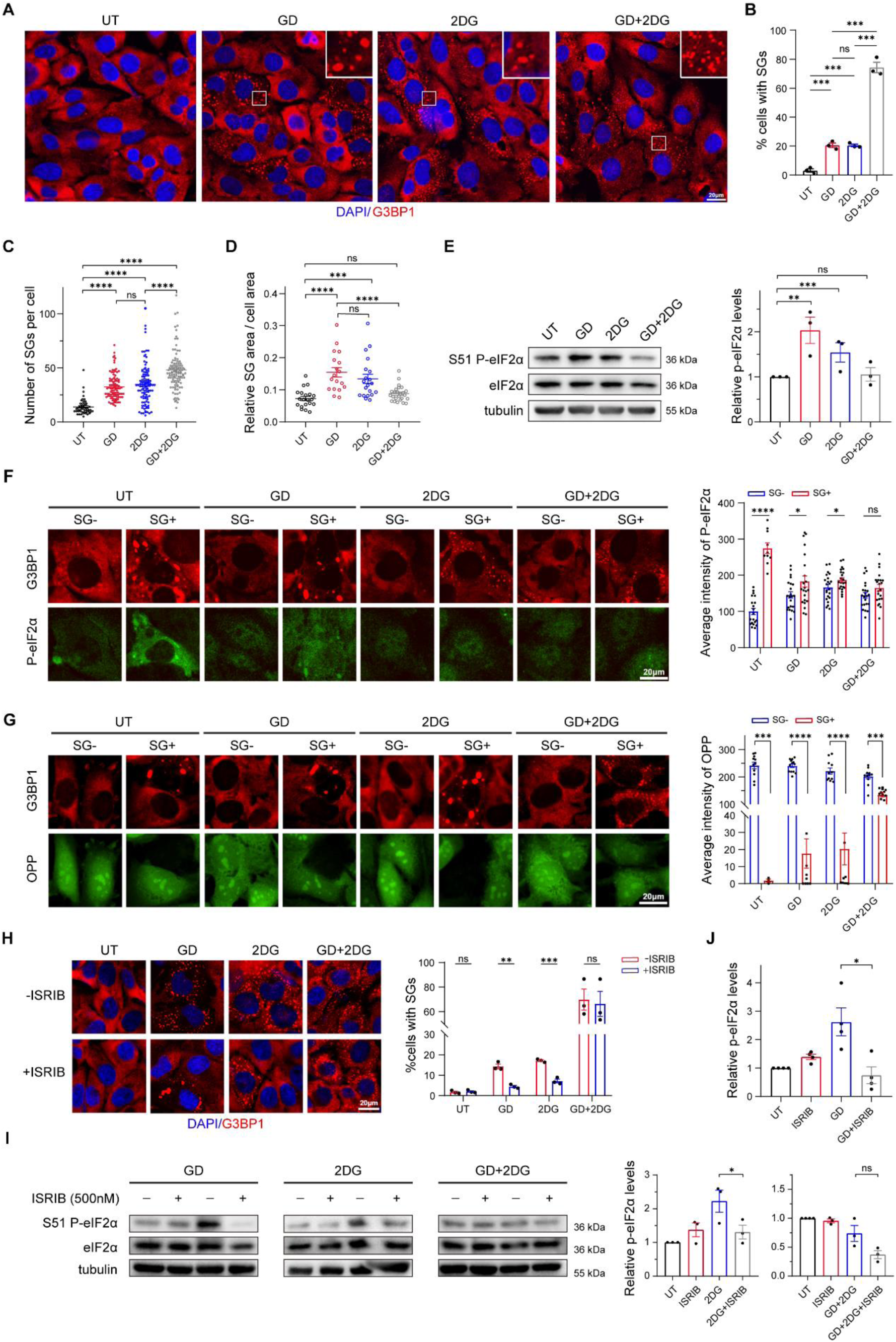
**SGs are differentially formed under different assaults against glycolysis.** (A) Representative images of SG formation (blue: DAPI, red: G3BP1) in 143B cells under no treatment (UT), glucose deprivation for 8 h (GD), treatment with 2DG for 1 h (2DG), and the combined treatment for 1 h (GD+2DG). Scale bar: 20 μm. Inset box: a zoom-in view of SGs. (B) Percentages of SG+ cells. Mean ± SEM (standard error of the mean) was calculated from three independent replicate experiments under each condition. Student’s t-test: **** *p* < 0.0001, *** *p* < 0.001, ** *p* < 0.01, * *p* < 0.05, ns. *p* > 0.05 (same throughout the manuscript). (C) Number of SGs per SG+ cell. For each condition, ≥ 49 individual cells were used for quantification. (D) Aggregated area proportion of SGs per SG+ cell. For each condition, ≥ 49 individual cells were used for quantification. (E) Western blot analysis of p-eIF2α (serine 51) and total eIF2α. Each condition has 3 independent replicate experiments. Quantification was performed as p-eIF2α/eIF2α ratio. (F) Representative images of fluorescent immunostaining (blue: DAPI, red: G3BP1, green: p-eIF2α) in SG- and SG+ cells under the four conditions, respectively. For each bar, average background-subtracted p-eIF2α intensity per unit cell area size was quantified from ≥ 10 individual cells. (G) Representative images of fluorescent immunostaining (blue: DAPI, red: G3BP1, green: OPP) in SG- and SG+ cells under the four conditions, respectively. For each bar, average background-subtracted OPP intensity per unit cell area size was quantified from ≥ 10 individual cells. (H) Representative images of SG formation (blue: DAPI, red: G3BP1) in cells under different glycolytic assaults combined with ISRIB. Each condition has 3 independent replicate experiments. (I-J) Western blot analysis and quantification of p-eIF2α and total eIF2α in cells under different glycolytic assaults combined with ISRIB. Each condition has 3 independent replicate experiments.

The pathway to SG formation can be either dependent or independent of phosphorylation of the translation initiation factor eIF2α (p-eIF2α). Western blot analysis revealed a significant increase of p-eIF2α/eIF2α ratio in both single-assault groups but not in GD+2DG (Figure 1E). We verified this phenomenon at the single-cell level through immunofluorescence staining. By comparing the signals of anti-p- eIF2α and O-propargyl-puromycin (OPP) between SG+ and SG- cells, we observed a correlation of SG presence to both p-eIF2α elevation and reduction in protein synthesis under single but not double assaults (Figure 1F-G). Consistently, GADD34, a factor involved in p-eIF2α dephosphorylation, was reduced under single but not double assaults (Figure S1). To verify the distinct dependences of p-eIF2α in SG formation in our system, we treated cells with ISRIB, an antagonist of ISR that acts downstream to all eIF2α kinases and specifically reverses the cellular effects of p- eIF2α (Sidrauski et al. 2015). Our results show that ISRIB effectively reduced both the fraction of SG+ cells and the level of p-eIF2α under single assaults, an effect that was diminished under GD+2DG (Figure 1H-J). Together, these results suggest that cells respond differently to a set of assaults that share the common target of glycolysis. For convenience, we refer to SGs formed under single and double assaults in our system as Type I and Type II SGs, respectively.

### OXPHOS gene transcripts are compartmentalized in Type II SGs

The protective role of SGs has been attributed to the compartmentalization of RNPs that can protect them from degradation and preserve their availability for future use if and when cells recover (Das et al. 2022). While the protein composition of SGs has been extensively studied, the specific mRNA species associated with SGs may also be informative about the specificity or diversity of responses to the inducing stress. To investigate the mRNA composition of SGs induced under our conditions, we generated an G3BP1-APEX2 fusion protein in a biotin-based proximity labeling experiment for enriching SG-associated mRNAs. Immunofluorescence staining showed subcellular colocalization between the fusion protein and the biotinylation signal (Figure 2A). We sequenced cDNA libraries prepared from bulk and G3BP1- associated mRNAs under no treatment (UT), GD, 2DG or GD+2DG at the peaking time for SG+ cells. Pairwise correlation between independent replicates (Pearson’s correlation *ρ* > 0.995) indicated an excellent data reproducibility. Importantly, only a minimal level of correlation (*ρ* = 0.172∼0.371) was observed between the bulk-seq and APEX-seq results within each pair (Figure 2B). These results support a selective enrichment of G3BP1-associated mRNAs under our experimental conditions.

**Figure 2.**
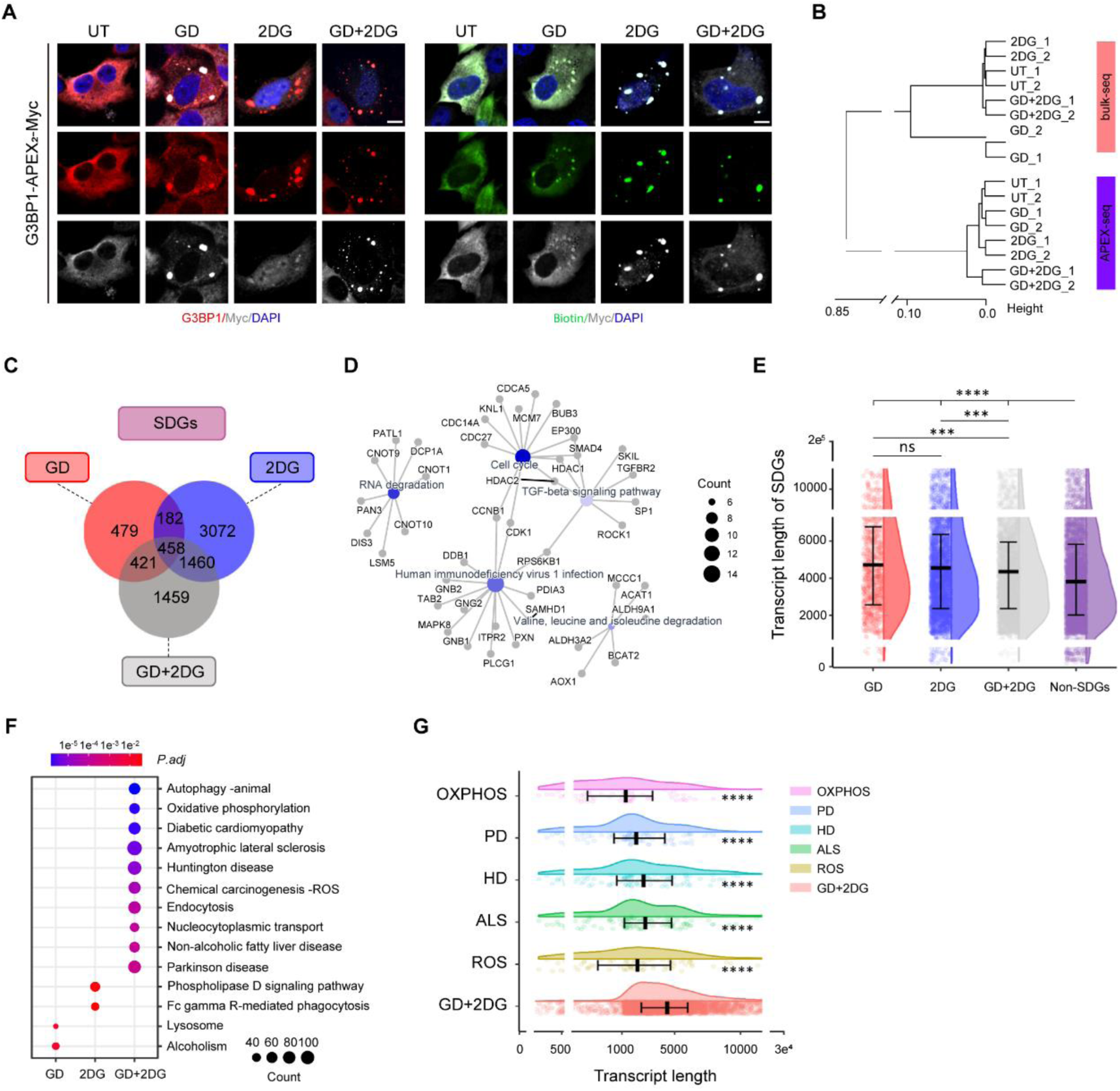
**Different glycolytic assaults result in different SG-associated transcripts.** (A) Colocalization of G3BP1-APEX2-Myc fusion proteins with biotinylation signals (blue: DAPI, red: G3BP1, gray: MYC, green: biotin) in cytoplasm and SGs under UT, GD, 2DG, and GD+2DG. Scale bar: 10 μm. (B) Hierarchical cluster analysis on pair-wise dissimilarities (calculated as 1 - Pearson’s correlation coefficient) among all bulk and G3BP1-APEX2-Myc-enriched RNA-seq datasets. Each condition has 2 independent replicates. (C) Venn diagram shows the numbers of stress-dependent G3BP1-associated genes (SDGs) resulted from GD, 2DG, and GD+2DG. (D) Gene pathways enriched with common SDGs among GD, 2DG, and GD+2DG. Blue dots represent enriched pathways, with darker colors indicating higher significance, and gray dots denote genes within specific pathway. (E) Transcript lengths of SDGs identified from GD, 2DG and GD+2DG and the other expressing genes (non-SDGs). Shown are the results based on the lengths of all quantified transcript isoforms. The trend remains consistent whether using transcript isoforms with the longest total length or the longest CDS (Figure S2). (F) Gene pathways enriched with SDGs identified from GD, 2DG and GD+2DG, respectively. (G) Transcript length distributions of SDGs in the GD+2DG-enriched pathways of OXPHOS, PD, HD, ALS, and ROS.

Based on a hierarchical clustering analysis, we found that, among all G3BP1- associated transcriptomes, the GD+2DG groups were most distant from the other groups (Figure 2B). To further evaluate the difference in the mRNA contents, we enriched G3BP1-associated transcriptomes by comparing each of the three stress conditions against those from UT. Such an analysis led to the identification of 1540, 5172 and 3798 stress-dependent, G3BP1-associated genes under GD, 2DG and GD+2DG, respectively (Figure 2C). We referred to these stress-dependent genes as SDGs. Among them, 458 genes were shared among all three conditions. Functional annotation analysis revealed that these common SDGs were enriched in “cell cycle”, “TGF-β signaling pathway”, “RNA degradation”, “valine, leucine and isoleucine degradation” and “human immunodeficiency virus 1 infection” (Figure 2D). These results, which confirm our recently reported effect of long-term glucose depletion on cell cycle progression (Zheng et al. 2023; see also Figure S4 for cell cycle analysis of the current study), are supportive of the hypothesis that SGs store away transcripts which encode proteins with “unwanted” functions at the time of stress.

Transcript length is one of the features that determine the selectivity of mRNAs sorting into SGs. Here we evaluated the transcript length distributions of SDGs identified from the three conditions of glycolytic stress. We found that, similar to endoplasmic reticulum (ER) stress or arsenite exposure (Namkoong et al. 2018; Khong et al. 2017; Ren et al. 2023; Van Treeck et al. 2018), all three glycolytic assaults resulted in SDGs whose transcripts were much longer than those of their non- SDG counterparts (Figure 2E). However, SDGs induced by GD+2DG had significantly shorter transcript lengths than those induced by GD or 2DG alone (Student’s t-test *p* = 0.004 and 0.03, respectively). This finer selection of SDG contents likely reflects a tailored response strategy to the double assault. Indeed, functional annotation analysis showed that SDGs identified from GD+2DG were significantly involved in neurodegenerative diseases such as amyotrophic lateral sclerosis (ALS), Huntington disease (HD) and Parkinson disease (PD; Figure 2F). Among the specifically-enriched pathways, genes involved in oxidative phosphorylation (OXPHOS) have the shortest transcript lengths, and many of these genes are shared by the pathways of ALS, HD, PD, and reactive oxygen species (ROS; Figure 2G). Together, our results suggest a qualitative difference in mRNA composition of SGs between single and double glycolytic assaults and, more importantly, a selective sequestration of OXPHOS gene transcripts by SGs induced by the double assault.

### Mitochondrial function is defective in cells that form Type II SGs

The preferential enrichment of OXPHOS genes in SGs formed under the double assault suggests a possibility of mitochondrial dysfunction. To test this, we first performed Western blot analysis on two selected proteins that are encoded by SDGs: NDUFB4 serves as an accessory subunit of NADH dehydrogenase (mitochondrial complex I), and COX6B1 constitutes a critical subunit of cytochrome c oxidase (complex IV). The levels of both proteins were significantly reduced under GD+2DG but not GD or 2DG alone (Figure 3A-B). To investigate whether the selective sequestration of the transcripts of mitochondrial SDGs led to a general mitochondrial dysfunction, we determined the levels of three other representative mitochondrial proteins not encoded by SDGs: TOM20 is a subunit of the outer mitochondrial membrane translocase (TOM complex), ND6 is a mitochondrial-encoded subunit of complex I, and ATP6 is a mitochondrial-encoded subunit of ATP synthase (complex V). Our results showed a significant reduction in the level of these mitochondrial proteins under GD+2DG but not GD or 2DG alone (Figure 3C-D). Consistently, GD+2DG treatment yielded a most prominent reduction in both TOM20 immunofluorescence staining signal and mitochondrial area size in SG+ cells relative to SG- cells (Figure 3E-F). Interestingly, we also detected a higher mitochondrial proximity of Type II SGs than those of SGs (Figure 3G), a phenomenon that has been reported to link SGs to metabolic remodeling (Amen et al. 2021). As expected of a mitochondrial dysfunction, the levels of intracellular ATP and mitochondrial respiration were significantly reduced under GD+2DG relative to GD or 2DG alone (Figure 3H-J). Together, these results document a unique feature of Type II SGs, a preferential retention of mitochondria-related mRNAs, and the accompanying mitochondrial defects.

**Figure 3.**
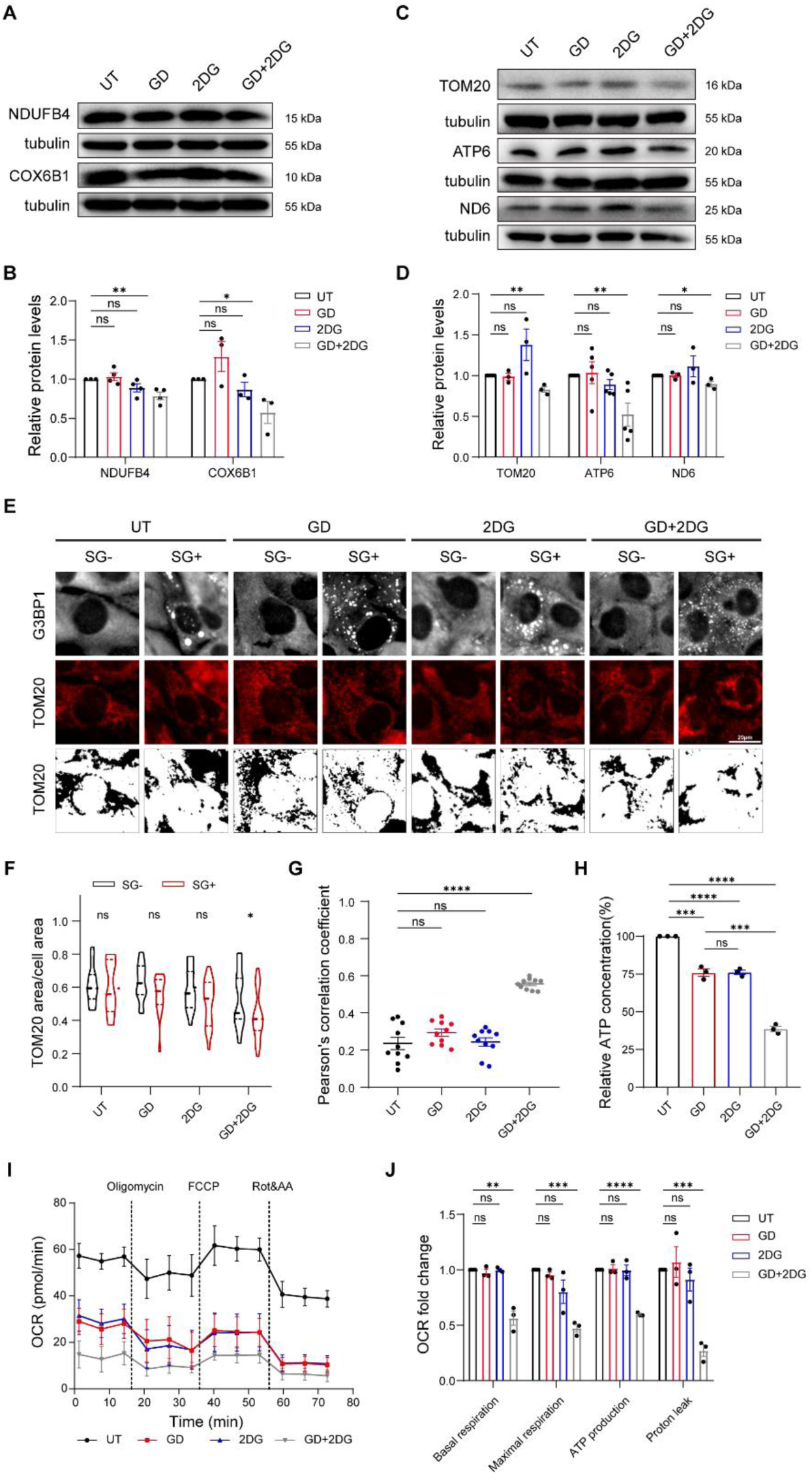
**Mitochondrial dysfunction under double assault against glycolysis.** (A-B) Western blot analysis of NDUFB4 and COX6B1, which were identified as SDGs under GD+2DG. (C-D) Western blot analysis of TOM20, ND6 and ATP6, which were identified as non-SDGs. (E-F) Representative images of immunofluorescence staining against G3BP1 (gray) and TOM20 (red and binary). Quantification was performed by normalizing the aggregated area size of identified mitochondria using TOM20 with the cell size. N ≥ 7 for each data point. (G) Pearson’s correlation coefficients show pixel intensity correlation between the two channels within single cells of panel E. The analysis was performed by “Colocalization Finder” in ImageJ. Each dot represents one single cell. (H) Relative levels of intracellular ATP under GD, 2DG and GD+2DG at the corresponding SG peaking times. (I) Oxygen consumption rate (OCR) was measured in cells sequentially treated with oligomycin, trifluoromethoxy carbonylcyanide phenylhydrazone (FCCP), and rotenone+antimycin A. Errorbars represent SEM computed from three independent measurements. (J) Relative rates of basal respiration, maximal respiration, ATP production and proton leak, normalized to the corresponding measurements in the side-by-side UT samples.

### Type II SGs differ from Type I SGs in their failure to dissolve during prolonged treatment

Previous studies suggest that intracellular ATP deficiency can prevent conventional SGs from disassembling (Eum et al. 2020; Jain et al. 2016; Wang et al. 2022). Based on our finding that mitochondrial defects were linked only to SGs formed under GD+2DG, we predicted that SGs formed under either GD or 2DG alone would engage in disassembly without the release of glycolytic insult, i.e., during prolonged treatment. Therefore, we analyzed the fate of SGs by extending the duration of treatments to 24 h. Under GD alone, the number of SG+ cells, the average SG number and the average SG size were all significantly reduced after 8 h (Figure 4A). At 24h, SG+ cells approached a minimum level (∼0.95%) that was basically a background level seen in untreated cells (∼1%). Similarly, under 2DG alone, these SG features continuously declined after the peaking time at 1 h (Figure 4B). In contrast, under GD+2DG, SGs were maintained even at 24 h in all examined cells (N = 1,091; Figure 4C). These differences in SG dynamics between GD or 2DG alone and GD+2DG were also observed in HeLa cells (Figure S5).

**Figure 4.**
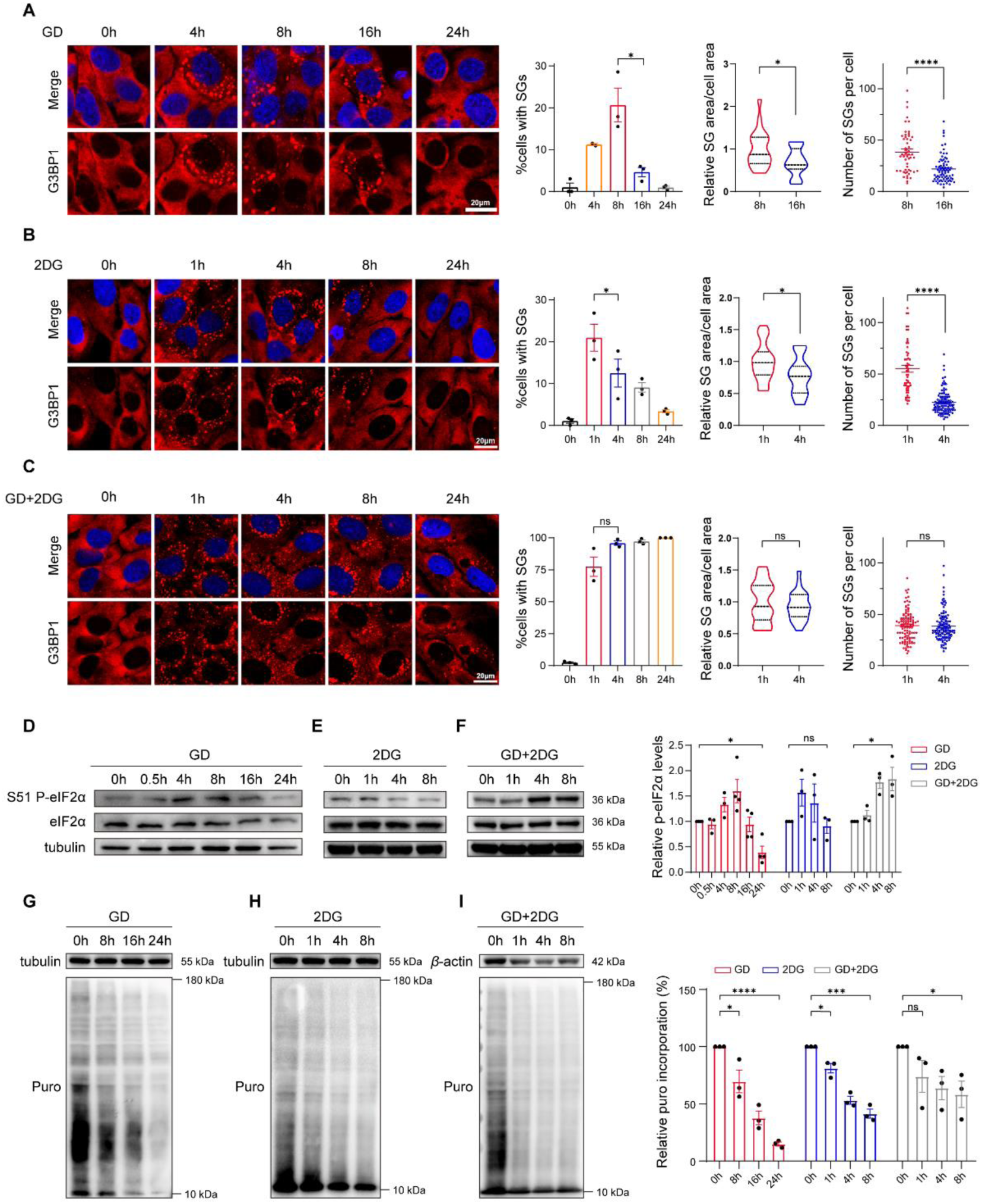
**Distinct stress granule dynamics of cells under different assaults against glycolysis.** (A-C) Representative images and quantifications of SG formation (blue: DAPI, red: G3BP1) in 143B cells treated by GD (A), 2DG (B), and GD+2DG (C) for different durations. (D-F) Western blot analysis of p-eIF2α (serine 51) and total eIF2α in cells treated by GD, 2DG, GD+2DG for different durations. Quantification shown on the right was performed by normalizing p-eIF2α to eIF2α. Errorbars represent SEM computed from three independent replicates. (G-I) Western blot analysis of puromycin incorporation assay in cells treated by GD, 2DG, GD+2DG for different durations. Quantification shown on the right was performed by normalizing each set of experiments to the control group at 0 h. Errorbars represent SEM computed from three independent replicates.

Given our result that eIF2α phosphorylation and translation repression are differentially impacted by double or single assaults, we sought to evaluate the dynamic properties of these processes during prolonged treatments. We detected a decline in p-eIF2α under GD or 2DG alone, which was correlated with SG dissociation dynamics (Figure 4D-E), whereas p-eIF2α exhibited a continuous increase under GD+2DG despite its insignificance at the time of SG formation (Figure 4F). Puromycin incorporation assays documented consistently continuous declines in global protein synthesis in all three conditions (Figure 4G-I), supporting that translation repression can be achieved through both p-eIF2α-dependent and - independent pathways. Together, these results document a dynamic disassembly process for Type I SGs formed under sustained single glycolytic assaults, a process lacking in cells that form Type II SGs.

### Cells forming Type II SGs upregulate the bulk mRNA expression of OXPHOS genes

To analyze transcriptional responses to different glycolytic assaults as a function of time, we generated bulk mRNA-seq datasets from cells under GD, 2DG or GD+2DG from different durations. We were particularly interested in comparing the two distinct phases of SG lifecycle: formation (0∼8, 0∼1 and 0∼1 h under GD, 2DG and GD+2DG, respectively) and dissolution/persistence (8∼16, 1∼4 and 1∼4 h under GD, 2DG and GD+2DG, respectively). We performed a time-series clustering analysis to identify gene ontologies (GOs) with specific dynamic patterns: rise-falling, fall-rising, monotonically rising, and monotonically falling (Figures 5A; see Materials and Methods). A total of 9 pathways shared the same dynamics in cells under all three treatments (Figure 5B), the majority of which (8/9) were monotonically rising such as “IRES dependent viral translational initiation” and “eukaryotic translation initiation factor 3 complex Eif3m” (Figure 5C). This result supports the known mechanism of IRES-mediated translation upon stress (Yang et al. 2019; Lacerda et al. 2019) as a common cellular response to different glycolytic assaults.

**Figure 5.**
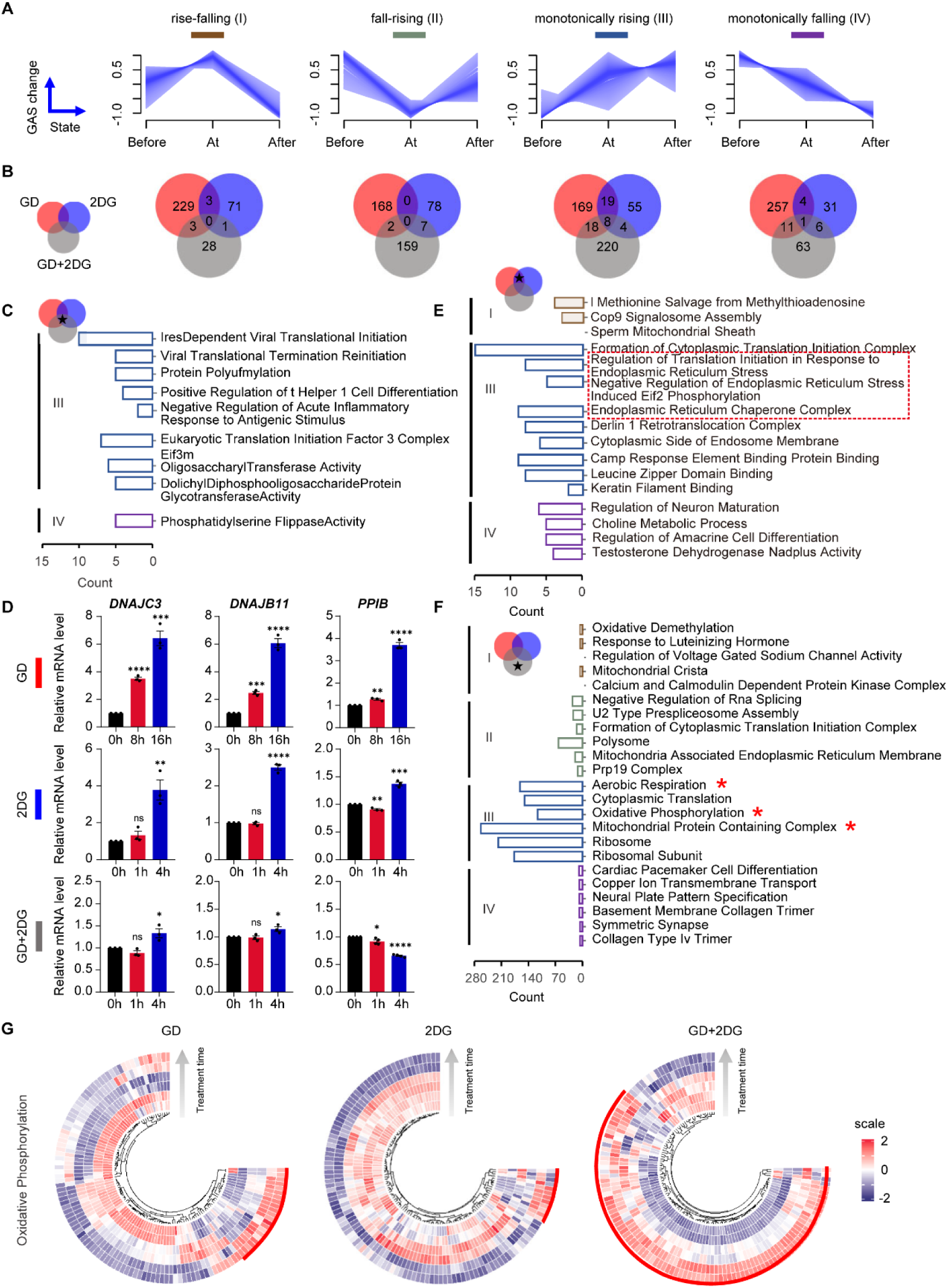
**Transcriptomic dynamics identifies upregulation of OXPHOS genes as a specific response to double glycolytic assaults.** (A) Geneset Activity Score (GAS) analysis identifies four clusters of gene ontologies (GOs) with distinct temporal patterns during the prolonged glycolytic assaults: rise- falling (I, dark brown), fall-rising (II, dark green), monotonically rising (III, dark blue), monotonically falling (IV, dark purple). X axis represents time points in relation to the SG peaking time; y axis represents changes in GAS, defined as the temporal variation in GSVA-derived scores of GO terms; each line represents one GO term. Shown is the result from GD; see Figure S3 for the results from 2DG and GD+2DG. (B) Venn diagrams illustrate the numbers of GOs with specific temporal patterns that are common or unique in cells under GD, 2DG and GD+2DG. (C) Gene pathways that share the same temporal pattern under all three conditions. Color codes are the same as in (A). (D) RT-qPCR analysis of *DNAJC3*, *DNAJB11* and *PPIB*, three genes identified in (E). Errorbars represent one standard deviation computed from 3 independent replicate experiments. (E) Gene pathways that share the same temporal pattern between GD alone and 2DG alone but not GD+2DG. Color codes are the same as in (A). (F) Gene pathways that exhibit a specific temporal pattern under GD+2DG but behave differently under GD or 2DG alone. Color codes are the same as in (A). (G) Heat maps showing scaled expression of OXPHOS genes during the prolonged treatments under GD, 2DG and GD+2DG.

There were 26 pathways that shared the same dynamics between GD alone and 2DG alone but not with GD+2DG. Among these, 3 monotonically-rising pathways were related to ER stress (Figure 5D-E), indicative of a transcription-level response to elevated p-eIF2α and reduced protein synthesis under either single assault. In contrast, among the pathways with a unique temporal pattern in cells under GD+2DG, the top significant GOs were related to mitochondrial functions, including “mitochondrial protein containing complex”, “aerobic respiration” and “oxidative phosphorylation” (Figure 5F). The majority of genes in these pathways exhibited an overall increasing trend as a function of treatment duration under GD+2DG but not under GD or 2DG alone (Figure 5G). It is particularly worth noting that this expression increase in OXPHOS genes coincided with the acute trapping of their transcripts by SGs (Figure 2F-G). Given the observation of mitochondrial defects including the reduced expression of OXPHOS genes, we postulate that cells were making a compensatory response to the double glycolytic assault, a response that would only end up being futile.

### Mitochondrial inhibition prevents Type I SG dissolution during prolonged 2DG treatment

To test whether mitochondrial defects under the double assault may underlie the inability of cells to reverse SGs, we analyzed cells under treatments of 2DG along with inhibitors that target different mitochondrial respiratory complexes. If our hypothesis is correct, we would expect mitochondrial inhibition to render SGs formed under Type I condition non-dissolvable. Our results show that, while each mitochondrial inhibitor alone showed little inducement of SGs, combining a mitochondrial inhibitor with 2DG led to a dramatic induction of SGs within 1 h (Figure 6A). In addition, both the fraction of SG+ cells and the p-eIF2α level were persistent during the prolonged treatments (Figure 6A-B). Furthermore, in ρ0 cells, which are depleted of mitochondrial DNA, GD alone induced SGs that shared similar dynamics with those formed in 143B cells under GD+2DG (Figure 6C-D).

**Figure 6.**
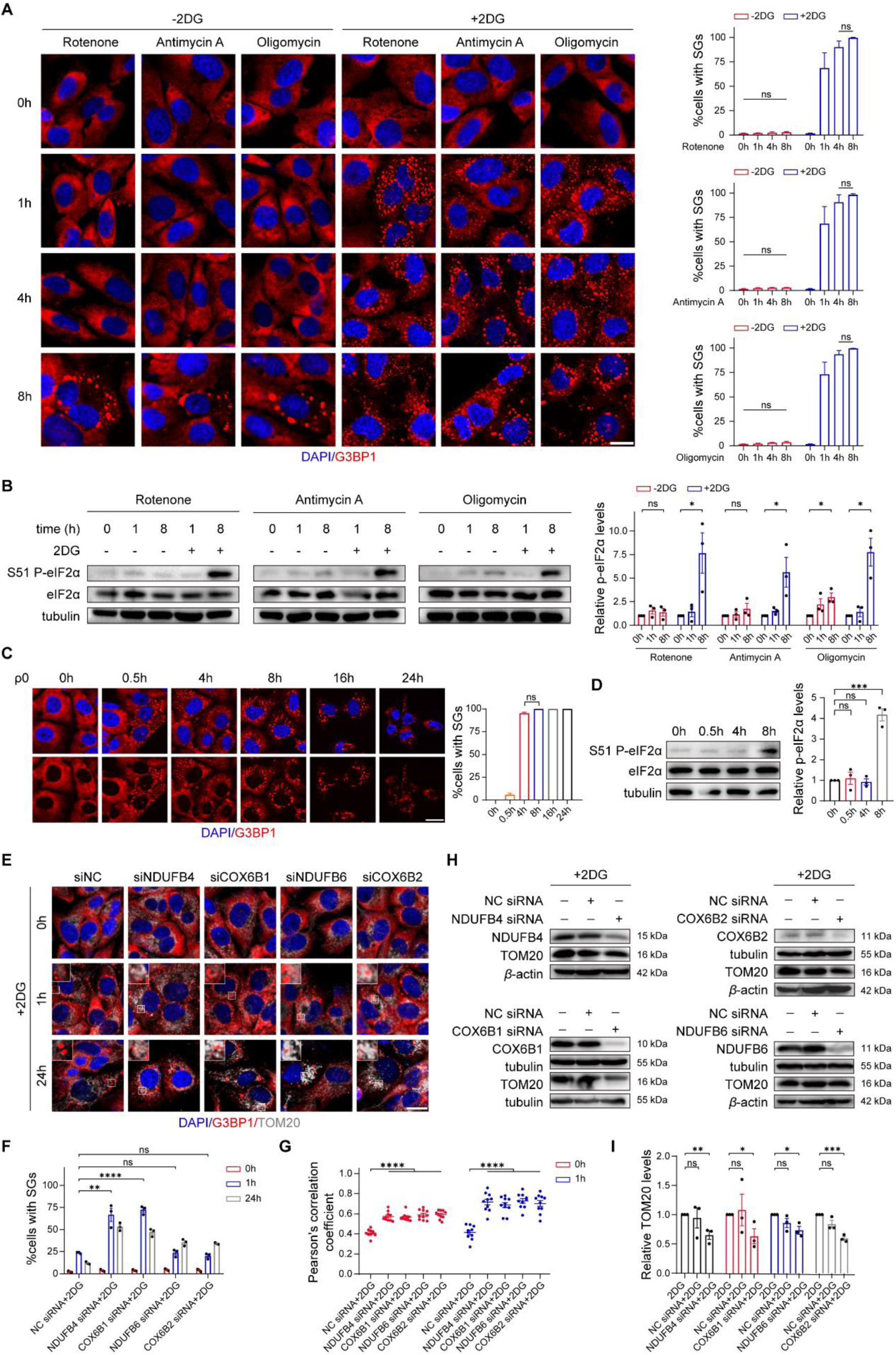
**Mitochondrial inhibition renders SGs formed under 2DG treatment non-dissociable.** (A) Representative images and quantifications of SG formation (blue: DAPI, red: G3BP1) in 143B cells treated with 2DG and mitochondrial inhibition (1 μM rotenone, 5 μM antimycin A, or 1.5 μM oligomycin). Scale bar: 20 μm (B) Western blot analysis of p-eIF2α (S51) and eIF2α in 143B cells treated with 2DG and mitochondrial inhibition. (C-D) Representative images and quantification of SG formation (blue: DAPI, red: G3BP1) and p-eIF2α in ρ0 cells were treated with GD for different durations. Scale bar: 20 μm. (E-F) Representative images and quantification of immunofluorescence staining against G3BP1 (red) and TOM20 (gray) in 143B cells treated with 2DG and a siRNA targeting negative control (NC), *NDUFB4*, *COX6B1*, *NDUFB6*, or *COX6B2*. Scale bar: 20 μm. (G) Pearson’s correlation coefficients quantify the spatial overlap between SGs (G3BP1 intensity) and mitochondria (TOM20 intensity) across treatments. Each dot represents measurements from one single cell. (H) Western blot analysis of TOM20, NDUFB4, NDUFB6, COX6B1, and COX6B2 proteins in 143B cells treated with or without 2DG and a siRNA. (I) Quantification of TOM20 protein levels in panel H.

To test whether a lack of any specific OXPHOS gene activity may lead to a similar phenotype, we combined siRNA knockdown with 2DG treatment. Here we tested two OXPHOS SDGs (*NDUFB4* and *COX6B1*) and two OXPHOS non-SDGs (*NDUFB6* and *COX6B2*). Figures 6E-I show that all of these siRNAs significantly decreased the protein level of TOM20 and, importantly, SGs formed under these conditions showed Type II hallmarks, including persistence at 24 h and proximal mitochondrial association (see also Figure 6E-F for a detectable difference in characteristics of SGs between SDG knockdown and non-SDG knockdown at 1 h). Together, these results support an involvement of mitochondrial activity in SG dissolution under prolonged glycolytic assaults, an engagement that, under the double assault, is diminished by SG sequestration of OXPHOS transcripts.

### Dissolution of Type I SGs requires HSP70 activity whose inhibition alters OXPHOS gene expression

Our results described thus far document a defining difference between single and double glycolytic assaults, i.e., the ability of SGs to dissociate under prolonged treatments (Figure 4A-C). To gain insights into the dissolution process of SGs formed under these assaults, we examined the potential dependence of SG clearance on autophagy and the HSP70 chaperone, respectively. We found that, while wortmannin, a potent autophagy inhibitor, affected neither SG formation nor dissolution (Figure 7A), siRNA against *HSPA1A* and VER-155008 targeting HSP70 both effectively preserved the fraction of SG+ cells during prolonged treatment of GD or 2DG alone (Figure 7B-C). Importantly, the expression level of HSP70 was significantly increased in cells undergoing SG dissolution but not in cells during prolonged GD+2DG treatment (Figure 7D). These results suggest a crucial role of HSP70 activity in SG dissolution during prolonged glycolytic stress.

**Figure 7.**
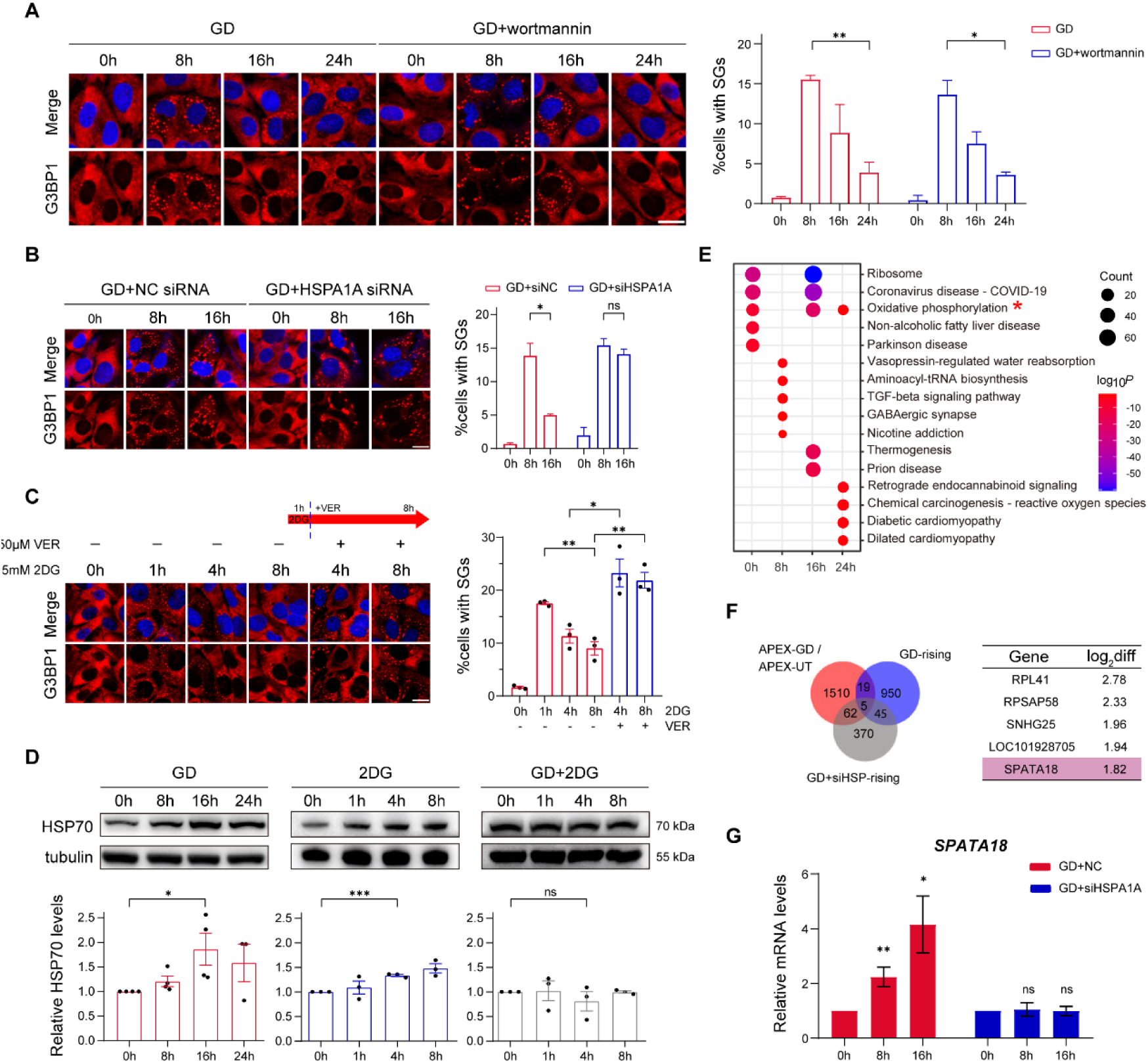
HSP70 is responsible for SG dissociation during prolonged treatments. (A-C) Representative images and quantifications of SG formation (blue: DAPI, red: G3BP1) in 143B cells treated with GD+ wortmannin (A), GD+*HSPA1A* siRNA (B) and 2DG+VER-155008 (C). Each experimental condition has at least 2 independent replicate experiments. (D) Western blot analysis of HSP70 in cells treated by GD, 2DG, GD+2DG for different durations. (E) Functional enrichment analysis of differential expression genes between GD and GD+siHSP at each corresponding time point. (F) Venn diagram shows the numbers of genes from SDGs under GD, upregulated at 16 h relative to 8 h under GD (GD-rising) and upregulated at 16 h relative to 8 h under GD+siHSP (GD+siHSP-rising). Left panel represents the top 5 genes from the intersection of SDGs under GD and HSP70-dependent SG-dissolution genes (upregulated at 16 h relative to 8 h under GD but not under GD+siHSP), ranked by the differences in upregulation levels between GD and GD+siHSP. (G) RT-qPCR analysis of *SPATA18* identified in (F). Errorbars represent the standard deviation calculated from 3 independent replicate experiments.

To explore HSP70-mediated regulation of SG dissolution during prolonged glycolytic stress, we performed bulk RNA-seq of cells that were preconditioned with *HSPA1A* siRNA and then co-treated with GD for different durations (GD+siHSP). Differential expression analysis between GD and GD+siHSP at each corresponding time point showed a significant enrichment of downregulated genes in OXPHOS at 0 and 16 h but not at 8 h (Figure 7E), suggesting a requirement of HSP70 activity for OXPHOS gene expression before the onset of glycolytic stress and during SG dissolution when the stress prolonged. In addition, by intersecting SDGs identified from G3BP1- APEX2 experiments and HSP70-dependent SG-dissolution genes (upregulated at 16 h relative to 8 h under GD but not under GD+siHSP), we obtained a list of 19 genes, which might be sequestered by SGs under the control of HSP70 (Figure 7F). Among these genes, *SPATA18* encodes the protein MIEAP, which promotes accumulation of lysosomal proteins in mitochondrial matrix and elimination of damaged proteins inside mitochondria (Ikari et al. 2024; Gaowa et al. 2018). RT-qPCR experiments confirmed that *SPATA18* was significantly increased from 8 to 16 h under GD only when HSP70 activity was intact (Figure 7G). Therefore, HSP70 may mediate the expression of mitochondrial quality control genes such as *SPATA18* to control SG dissolution.

### Cell survival under prolonged glycolytic assaults is correlated with SG dissolution

To test whether SG dissolution is an adaptive function that supports cell survival during prolonged glycolytic assaults, we performed flow cytometry with propidium iodide and Annexin V staining. We detected a pronounced fraction of apoptotic cells at 24 h under GD+2DG, but not under GD or 2DG alone (Figure 8A). Consistently, genes involved in apoptosis were enriched in upregulated genes under prolonged GD+2DG but not under prolonged GD or 2DG alone according to bulk mRNA-seq and RT-qPCR experiments (Figure 8B-C). Furthermore, when the dissolution of Type I SGs was blocked by HSP70 inhibition (GD+VER or 2DG+VER), the fraction of apoptotic cells was significantly increased (Figure 8D). Together these results show that cell survival is correlated with SG dissociation under prolonged glycolytic assaults, and that persistent SGs represent a sign for cells reaching an impasse.

**Figure 8.**
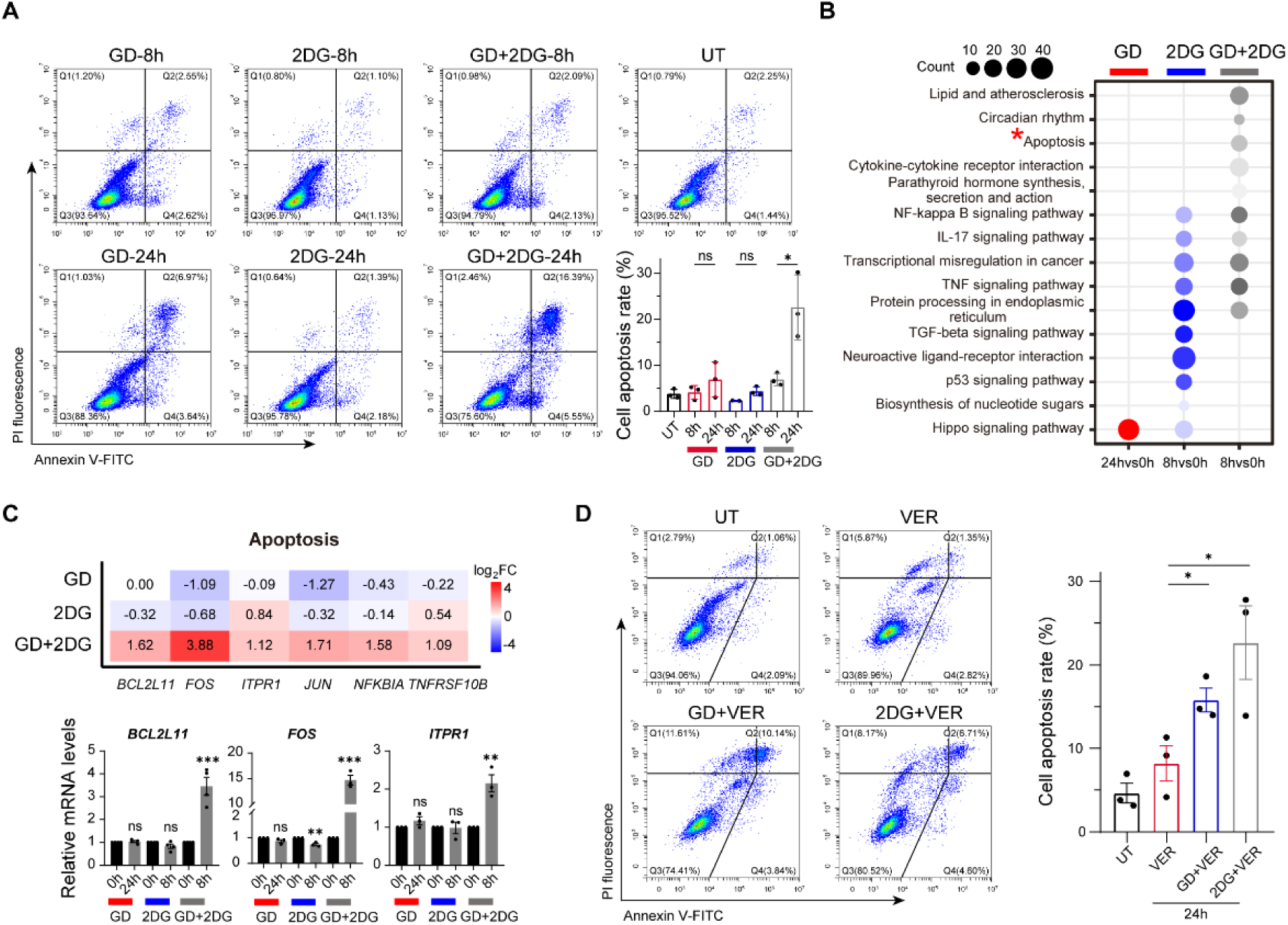
**Cell survival under prolonged glycolytic assaults.** (A) Flow cytometry analysis of apoptosis using Annexin V and propidium iodide staining in cells under UT, GD, 2DG, and GD+2DG for 8 or 24h. Errorbars represent SEM computed from three independent replicates. (B) Gene pathways enriched with upregulated genes under prolonged GD (24h), 2DG (8h) and GD+2DG (8h), respectively. (C) Heatmap for fold changes of genes involved in apoptosis (resulted from (B)) under prolonged GD, 2DG and GD+2DG, compared to the control. RT-qPCR confirmed the increase in the mRNA levels of *BCL2L11*, *FOS* and *ITPR1*, three genes in the apoptosis pathway, under GD+2DG but not under GD or 2DG alone. (D) Flow cytometry analysis of apoptosis in cells treated with VER-155008, GD+VER, and 2DG+VER. Errorbars represent SEM computed from three independent replicates.

## Discussion

In this study, we document a divergent formation of two distinct types of SGs under glycolytic inhibition. Type I SGs, which are induced by single assaults (GD or 2DG), resemble arsenite-induced SGs in their dependence on eIF2α phosphorylation for assembly and HSP70 activity for disassembly. In contrast, Type II SGs, which are induced by GD+2DG and previously referred to as energy deficiency-induced SGs (eSGs), exhibit a distinct biochemical state characterized by the sequestration of OXPHOS gene transcripts and mitochondrial dysfunction. Our comparative analysis of these two types of SGs suggests a feedback loop between the formation of Type II SGs and mitochondrial dysfunction, a regulatory loop that can render Type II SGs non-dissociable under sustained stress (see Figure 9 for a graphic model). Effectively the sequestration of OXPHOS transcripts within SGs under double assault widens the impact of glycolytic inhibition to further exacerbate energy deficit and prevent SG disassembly. This feedback loop highlights the dual roles of SGs in stress adaptation and pathogenesis, providing insights into the molecular mechanisms underlying SG- associated diseases.

**Figure 9.**
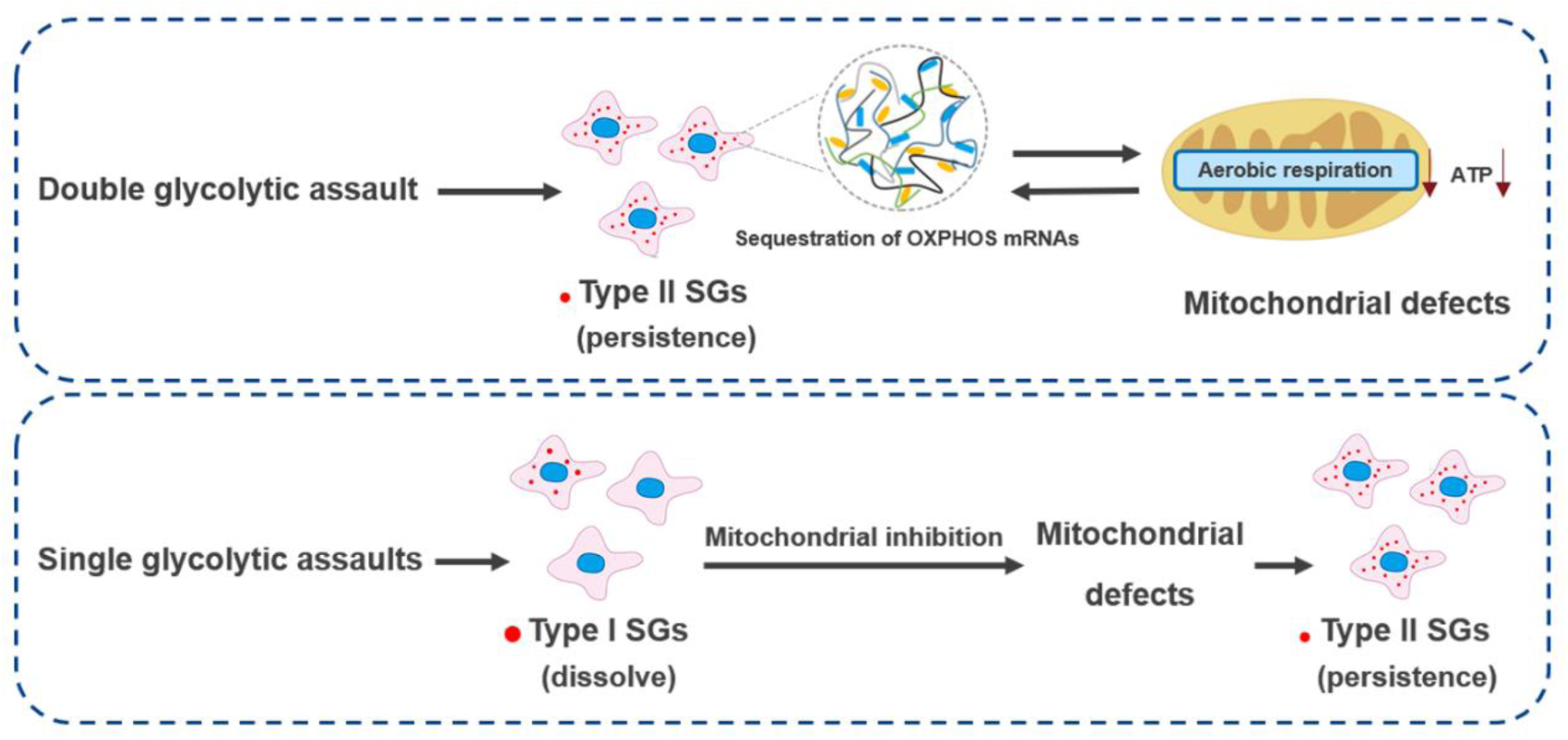
**A model showing the formation of two distinct types of SGs under glycolytic inhibition.** The double assault (glucose depletion + 2DG treatment) leads to the generation of Type II SGs, which specifically sequester mRNA molecules related to oxidative phosphorylation and intensify cellular energy deficits. This energy crisis drives Type II SGs into a non-dissolvable state. By contrast, single glycolytic assaults lead to the formation of Type I SGs, which can naturally dissolve during prolonged treatment. Type I SGs may be transformed into the non-dissolvable state by additional mitochondrial inhibition (or HSP70 dysfunction). This model depicts the scenario where SGs serve as both a stress responder and a driver of metabolic collapse, offering mechanistic insight into the association of pathological SGs.

The dynamic and reversible nature of physiological SGs enables cells to adapt to and recover from stressful conditions. However, the persistence of SGs containing pathological contents can lead to cell dysfunction and death (Ivanov et al. 2019; Zhang et al. 2019; Mahboubi et al. 2017; Sato et al. 2024). Such a paradoxical divergence highlights a delicate control of the quality and dynamics of SGs. In fact, disassembly upon the release of the inducing stressors such as sodium arsenite and heat shock has been defined as a key feature of acute SGs (Marmor-Kollet et al. 2020; Hofmann et al. 2021; Buchan et al. 2009). Interestingly, SGs that are induced by the proteasome inhibitor MG132 can also disassemble during a long-term treatment without eliminating the stressor (Ganassi et al. 2016; Mazroui et al. 2007; Wang et al. 2022). Similarly, our Type I SGs, but not Type II SGs, are dissolved within 24 h when glycolytic restriction remains in effect. Therefore, different regimens of glycolytic stress induce distinct SG responses, resembling a physiological-to-pathological transition that accompanies disease progression. In this context, it is worth noting that, while the initiating differences between the single and double assaults on glycolytic inhibition might not be major on their own, the positive feedback loop that involves the sequestration of OXPHOS mRNAs and the persistent nature of type II SGs likely have contributed significantly to the bifurcation in the SG types.

It has been shown that severe energy deficiency can prevent arsenite-induced SGs from disassembling (Jain et al. 2016; Wang et al. 2022). Our time-resolved transcriptomic analysis shows that the bulk mRNA level of OXPHOS genes exhibits an increase along the GD+2DG treatment time, but this increase in mRNA expression is ultimately unsuccessful because their proteins remain at a reduced level and mitochondrial functions remain impaired. We suggest that the preferential sequestration of OXPHOS gene transcripts by Type II SGs is responsible for undercutting the effect of the compensatory transcriptional upregulation, leading to a perpetual mitochondrial dysfunction and energy stress, accompanied by a persistence of SGs and increased cell death.

There is a long-standing hypothesis that SGs serve as temporal storages and silent sites of untranslated mRNAs and unused RNA-binding proteins (Kedersha et al. 2002; Ivanov et al. 2019). Our APEX2-based transcriptomic analysis uncovers distinct profiles of G3BP1-associated transcripts between Type I and Type II SGs. Notably, Type II SGs preferentially sequester shorter transcripts, including OXPHOS genes, despite the conventional preference for long transcripts due to enhanced RNA-RNA interactions (Campos-Melo et al. 2021; Ren et al. 2023; Lee et al. 2019; Khong et al. 2017). This unique mRNA composition reflects a specific cellular response to complete glycolytic inhibition and further underscores the role of Type II SGs in modulating mitochondrial function.

In conclusion, our work suggests a mechanism through which SG formation can transition between their protective and deleterious roles. Such a transition can be achieved through the operation of a feedback loop between SG dynamics and mitochondrial function. Our study provides a fresh perspective for understanding the pathogenesis of SG-associated diseases and potential therapeutic targets.

## Materials and Methods

### Cell culture and treatment

143B and HeLa cells were cultured in DMEM (Gibco) with 10% FBS, 100 U/ml penicillin and 100 U/ml streptomycin under 5% CO₂ at 37°C. The mitochondrial DNA-less ρ⁰ 206 cells, derived from 143B cells, were cultured under the same condition except an addition of 50 μg/ml uridine. For glycolysis inhibition, cells were rinsed in PBS and then transferred to glucose-free DMEM, DMEM with 25 mM 2- deoxy-D-glucose (2DG), or glucose-free DMEM with 2DG for various durations as described in main text. For mitochondrial inhibition, cells were rinsed in PBS and then transferred to DMEM with 1 μM rotenone, 5 μM antimycin A or 1.5 μM oligomycin. For p-eIF2α inhibition, cells were rinsed in PBS and then transferred to DMEM with 500 nM ISRIB (MCE, HY-12495A) for 1 h. For autophagy inhibition, cells were rinsed in PBS and then transferred to DMEM with 1 μM wortmannin (MCE, HY-10197) for 8 h. For HSP70 inhibition, cells were rinsed in PBS and then transferred to DMEM with 50 μM VER-155008 (MCE, HY-10941).

### siRNA transfection

All siRNAs were designed by DSIR (http://biodev.extra.cea.fr/DSIR/DSIR.html) and synthesized by GenePharma. For transfection, cells were treated with an siRNA at a final concentration of 50 nM using jetPRIME (Polyplus) for 36 h before other treatments. The oligo sequences are listed in Table S1.

### Quantitative RT-PCR

Cells were seeded in 6-well plates to reach approximately 80∼90% confluence, and total RNA was extracted using TRIzol (TaKaRa, 9109). The purified RNA was quantified by NanoDrop Spectrophotometer, and 1 μg RNA was subjected to reverse transcription using ABScript III RT Master Mix with gDNA Remover (ABclonal, RK20429). Quantitative PCR was performed using 2× Universal SYBR Green Fast qPCR Mix (ABclonal, RK21203). For each experimental group, three independent reverse transcription experiments were conducted and β-actin was used as an internal control. The primers are listed in Table S1.

### Western blot

Cells were washed with PBS and then lysed in RIPA buffer (FUDE, FD009) with Benzonase Nuclease (Beyotime, D7121) and complete Protease and phosphatase inhibitor cocktail (Beyotime, P1048). Lysates were loaded onto 10% SDS-PAGE and proteins were transferred to PVDF membranes (Millipore, IPVH00010). Membranes were incubated with rocking first in TBST with 5% milk at room temperature for 1 h, then in TBST with 5% milk and primary antibody at 4°C overnight. After three washes, membranes were incubated in TBST with secondary antibody and 5% milk at room temperature for 1 h. After another three washes, ECL Western Blotting Substrate (Vazyme Biotech, E412-01) were used for detection.

The following primary antibodies were used: HRP-conjugated β-Actin Rabbit mAb (1:5000, ABclonal, AC028), HRP-conjugated β-Tubulin Mouse mAb (1:5000, ABclonal, AC030), rabbit anti LC3B (1:1000, ABclonal, A19665), mouse anti HSP70/HSPA1 (1:2000, ABclonal, A1507), rabbit anti HSP70 (1:2000, ABclonal, A23457), rabbit anti phospho-eIF2α (S51; 1:1000, Cell Signaling, 3398), mouse anti phospho-EIF2S1 (Ser51) (1:1000, proteintech, 68023-1-Ig), rabbit anti EIF2S1 (1:1000, proteintech, 82936-1-RR), rabbit anti EIF2S1/ EIF2A (1:1000, proteintech, 11170-1-AP), rabbit anti GADD34 (1:1000, proteintech, 10449-1-AP), mouse anti TOM20 (1:1000, proteintech, 66777-1-Ig), rabbit anti TOM20 (1:1000, proteintech, 11802-1-AP), rabbit anti ATP6 (1:1000, ABclonal, A23150), rabbit anti COX6B1 (1:1000, proteintech, 11425-1-AP), rabbit anti MT-ND6 (1:1000, ABclonal, A17991), rabbit anti NDUFB4 (1:1000, proteintech, 27931-1-AP), rabbit anti COX6B2 (1:1000, proteintech, 11437-1-AP), rabbit anti NDUFB6 (1:1000, proteintech, 16037-1-AP).

### Immunofluorescence staining

Cells were washed in PBS, fixed in 4% paraformaldehyde at room temperature for 20 min, and rinsed in PBS buffer containing 0.1% Triton X-100 (PBST) for 10 min. Then cells were blocked in PBST with 3% BSA for 1 h and incubated at 4°C overnight with primary antibody. After three washes, cells were incubated in PBST with secondary antibody and 3% BSA at room temperature for 1 h. After another three washes, cells were mounted in Antifade Mounting Medium with DAPI (Beyotime, P0131).

The following primary antibodies were used: rabbit anti G3BP1 (1:200, ABclonal, A3968), mouse anti TOM20 (1:200, proteintech, 66777-1-Ig), rabbit anti TOM20 (1:200, proteintech, 11802-1-AP), 488-conjugated G3BP1 pAb (1:200, proteintech, CL488-13057), mouse anti p-EIF2S1 (Ser51) (1:200, proteintech, 68023-1-Ig), rabbit anti p-eIF2α (1:200, Cell Signaling, 3398), mouse anti Myc tag (1:500, proteintech, 60003-2-Ig).

### OPP staining

O-propargyl puromycin (OPP) staining was performed using Click-iT™ Plus Alexa Fluor™ 555 Picolyl Azide Toolkit (Thermo Fisher, C10642) according to the manufacturer’s instruction. After three washes, cells were mounted in Antifade Mounting Medium with DAPI (Beyotime, P0131).

### Image analysis and quantification

For each glass slide, >= 5 different fields were imaged by Olympus FV1000 Confocal Microscope or Nikon Instruments A1 Confocal Laser Microscope. ImageJ tools were used to identify individual cells (based on DAPI signals) and quantify SG characteristics (based on G3BP1 signals). Cells with at least five G3BP1-positive spots detected in the cytoplasm were considered as SG+. In each SG+ cell, the number of SGs was measured using Analyze Particles and the aggregated area size of SGs was measured using ROI Manager.

For quantification of p-eIF2α and OPP, the fluorescence intensities within each identified cell were summed, background-subtracted and normalized to the cell area size using ImageJ. For quantification of TOM20, the total area size of fluorescent signals was measured by ROI Manager and normalized to the cell area.

### RNA sequencing

Total RNA was extracted using RNAiso Plus (TaKaRa, 9109). The libraries were generated using VAHTS Universal V8 RNA-seq Library Prep Kit for Illumina (Vazyme, NR605). Sequencing was performed on Novaseq 6000 (Nanjing Jiangbei New Area Biopharmaceutical Public Service Platform). Read quality was assessed using fastqc and adaptor sequences were removed using trim_galore v0.6.10. Then reads were aligned to GRCh38 using hisat2 v2.2.1, and summarized using featureCounts v2.0.1. Quantification of transcript isoforms was performed using StringTie v2.2.1 for assembly and analysis. Differential expression analysis was conducted using edgeR v4.4.0 with a cutoff fold change > 2 and adjusted *p*-value < 0.05. Functional annotation, including Gene Ontology (GO) and Kyoto Encyclopedia of Genes and Genomes (KEGG) enrichment analyses, was carried out using clusterProfiler v4.14.3 with adjusted *p*-value < 0.05.

### APEX-based proximity labeling and RNA sequencing analysis

G3BP1-APEX2-Myc was generated by an in-frame fusion of G3BP1, APEX2 and Myc tag in pAcGFP1-N1 vector. After transfection, cells were cultured for 48 h to have adequate expression, and then subjected to different experimental treatments. The resulting samples were biotin-labeled as previously described (Somasekharan et al. 2020). For immunofluorescence staining analysis, rabbit anti G3BP1 (1:200, ABclonal, A3968), mouse anti Myc tag (1:500, proteintech, 60003-2-Ig) and Alexa Fluor™ 647-conjugated streptavidin (1:400, Thermo Fisher, S21374) antibodies was used. For RNA-seq analysis, biotinylated RNAs were pulled down using C1 Streptavidin beads (Thermo Fisher) according to the manufacturer’s instruction. To identify stress-dependent G3BP1-associated genes (SDGs), we performed differential expression analysis by comparing APEX-seq under a given stress condition over APEX-seq under no treatment in edgeR v4.4.0 with a cutoff fold change > 2 and adjusted *p*-value < 0.05.

### Time-series RNA sequencing analysis

We generated bulk mRNA-seq datasets from cells before, at and after the SG peaking time under GD (0, 8 and 16 h), 2DG (0, 1 and 4 h) or GD+2DG (0, 1 and 4 h). For each dataset, we used GSVA v2.0.1 to calculate Geneset Activity Score (GAS) of all GO terms, limma v3.62.1 to identify GO terms with differential GASs (fold change > 2 and adjusted *p*-value < 0.05), and Mfuzz v2.66.0 to cluster differential GOs with the same dynamic patterns of GAS.

### FACS analysis of ROS, apoptosis and cell cycle

Cells were treated with Reactive Oxygen Species Assay Kit (Beyotime, S0035M), Annexin V-FITC assay kit (Beyotime, C1062) or Cell Cycle and Apoptosis Analysis Kit (Beyotime, C1052) according to the manufacturer’s instructions, respectively, and then fluorescence was quantified by flow cytometry.

### Measurement of ATP

Cells were assayed by Enhanced ATP Assay Kit (Beyotime, S0027) according to the manufacturer’s instruction.

### Measurement of Oxygen Consumption rate

Cells were plated onto a Seahorse XF96 Cell Culture Microplate (Agilent) at a density of 1.0 × 10^4^ cells/well. After an overnight incubation with 5% CO₂ at 37 °C, the culture medium was replaced by the assay medium containing 1 mM pyruvate, 4 mM glutamine, and with or without 25 mM glucose (as the control and the GD group respectively). Then sequential treatments with 1 µM oligomycin, 2 µM FCCP, and 1 µM rotenone+antimycin A allowed for generating the full profile of OCR.

### Statistics

Each sequencing data has at least two replicate samples. Each quantitative experiment has at least three independent samples. Unless otherwise stated in the legend, all quantitative results were presented as mean ± standard error of the mean (SEM), and analyses of the mean were presented as unpaired two-tailed Student’s t- test.

## Data availability

All raw RNA-seq data generated in this study have been submitted to the NCBI BioProject database under accession number PRJNA1289020.

## Acknowledgements

This study was supported by the National Natural Science Foundation of China (32470584) and the National Key R&D Program of China (2021YFC2700403). We acknowledge support of Zhejiang University School of Medicine affiliated Women’s Hospital.

## Author contributions

W.Z., J.M. and F.H. conceived the study and designed the experiments; W.Z., M.X. and X.L. performed experiments and generated data; W.Z. and R.X. analyzed the data and generated all figures; Y.G. and M.G. provided technical and managerial support; J.M. and F.H. acquired funding; W.Z., R.X., J.M. and F.H. wrote the paper and all approved the paper.

---

**Figure S1.**
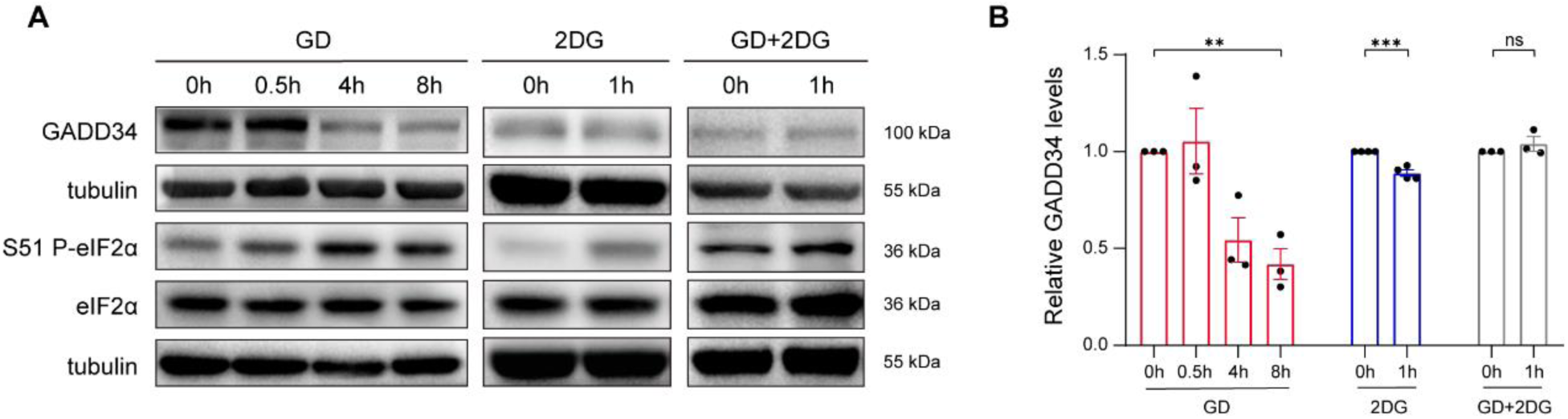
**GADD34 is reduced and negatively correlated with p-eIF2*α* under single glycolytic assaults but not under double assault.** (A-B) Western blot analysis of GADD34, p-eIF2α, and eIF2α in cells under GD, 2DG or GD+2DG. Quantification was performed by normalizing p-eIF2α to eIF2α and normalizing GADD34 to its corresponding internal control. Mean ± SEM was calculated from three independent replicate experiments.

**Figure S2.**
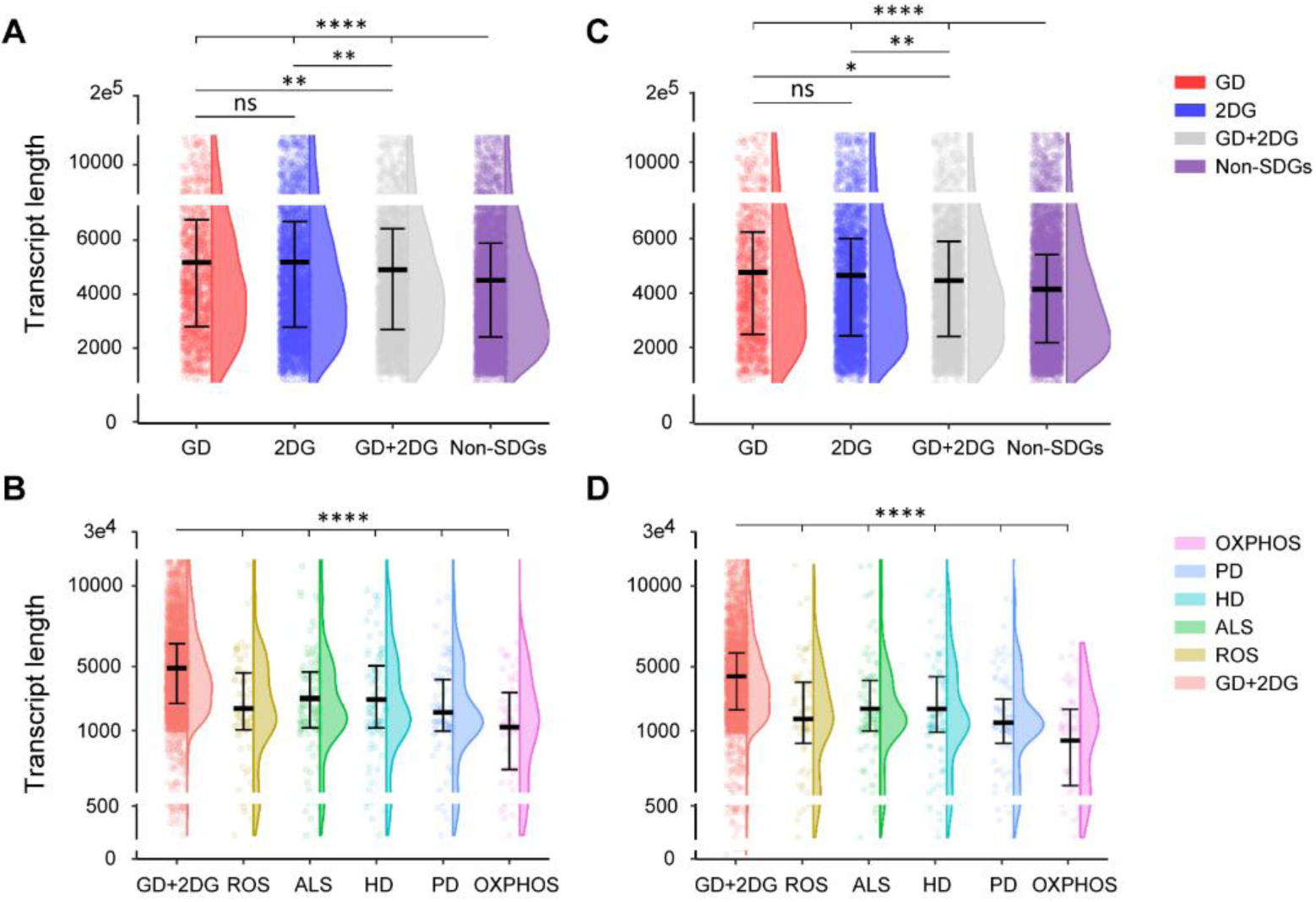
**Comparison of transcript lengths under different conditions.** (A, C): Transcript lengths of SDGs resulting from GD, 2DG, GD+2DG and non- SDGs, measured using the longest total length (A), or the longest CDS (C), respectively. (B, D): Distribution of transcript lengths for SDGs enriched in specific pathways under GD+2DG, including OXPHOS, PD, HD, ALS, and ROS, measured using the longest total length (B) or the longest CDS (D), respectively.

**Figure S3.**
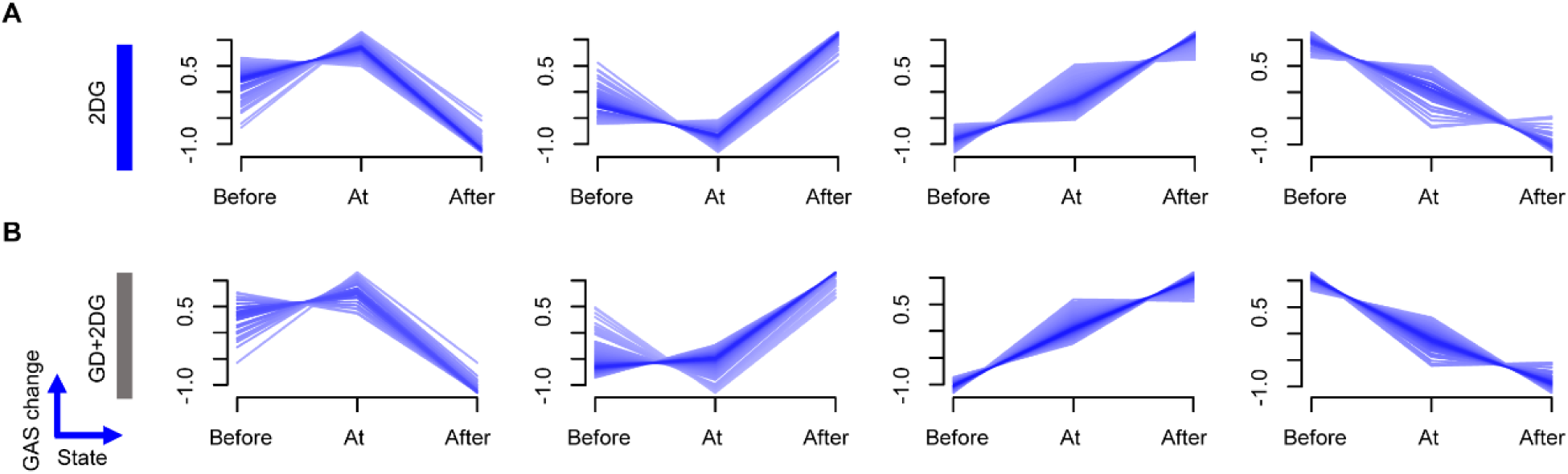
**Time-course Geneset Activity Score (GAS) analysis under different conditions.** (A-B) Same as Figure 5A but results from 2DG (A) and GD+2DG (B), respectively.

**Figure S4.**
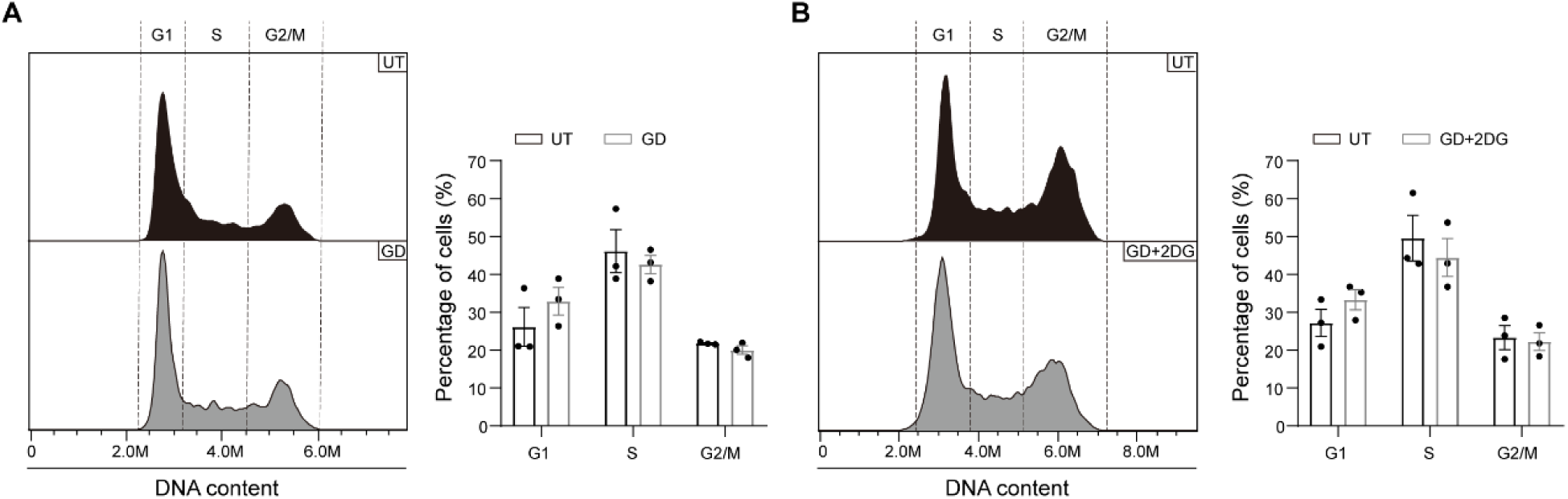
**Cell cycle is similarly disturbed by different glycolytic assaults.** (A-B) Flow cytometry analysis of cell cycle using PI staining in cells under GD for 8h (A) and GD+2DG for 1h (B). Both treatments present a moderate increase in the proportion of G1 cells: from 28.63 ± 7.72% to 32.61 ± 6.29% under GD and from 27.12 ± 6.21% to 32.32 ± 4.41% under GD+2DG.

**Figure S5.**
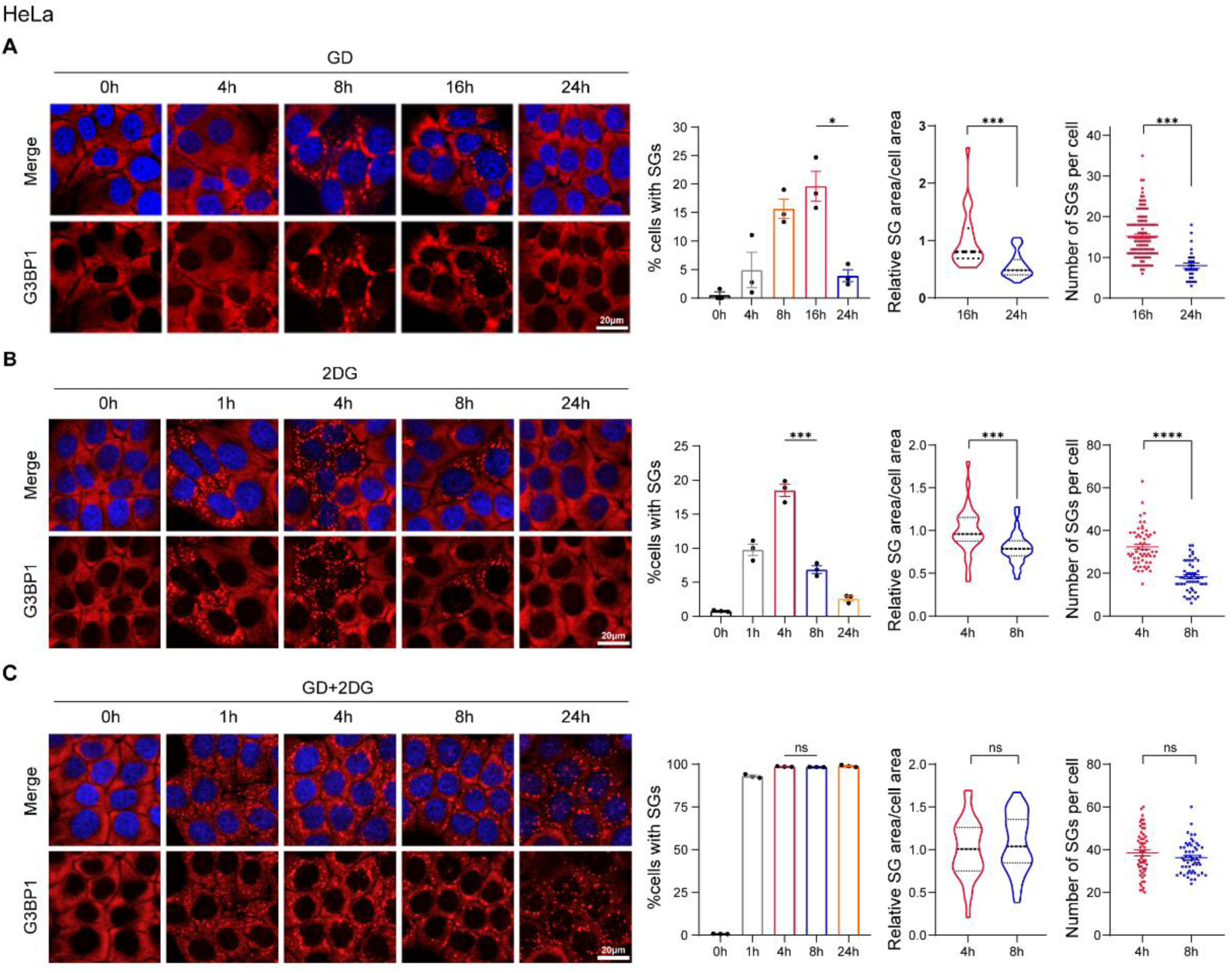
**Distinct stress granule responses of HeLa cells under different glycolytic assaults.** (A-C) Same as Figure 2A-C but results from HeLa cells.

**Table S1.**
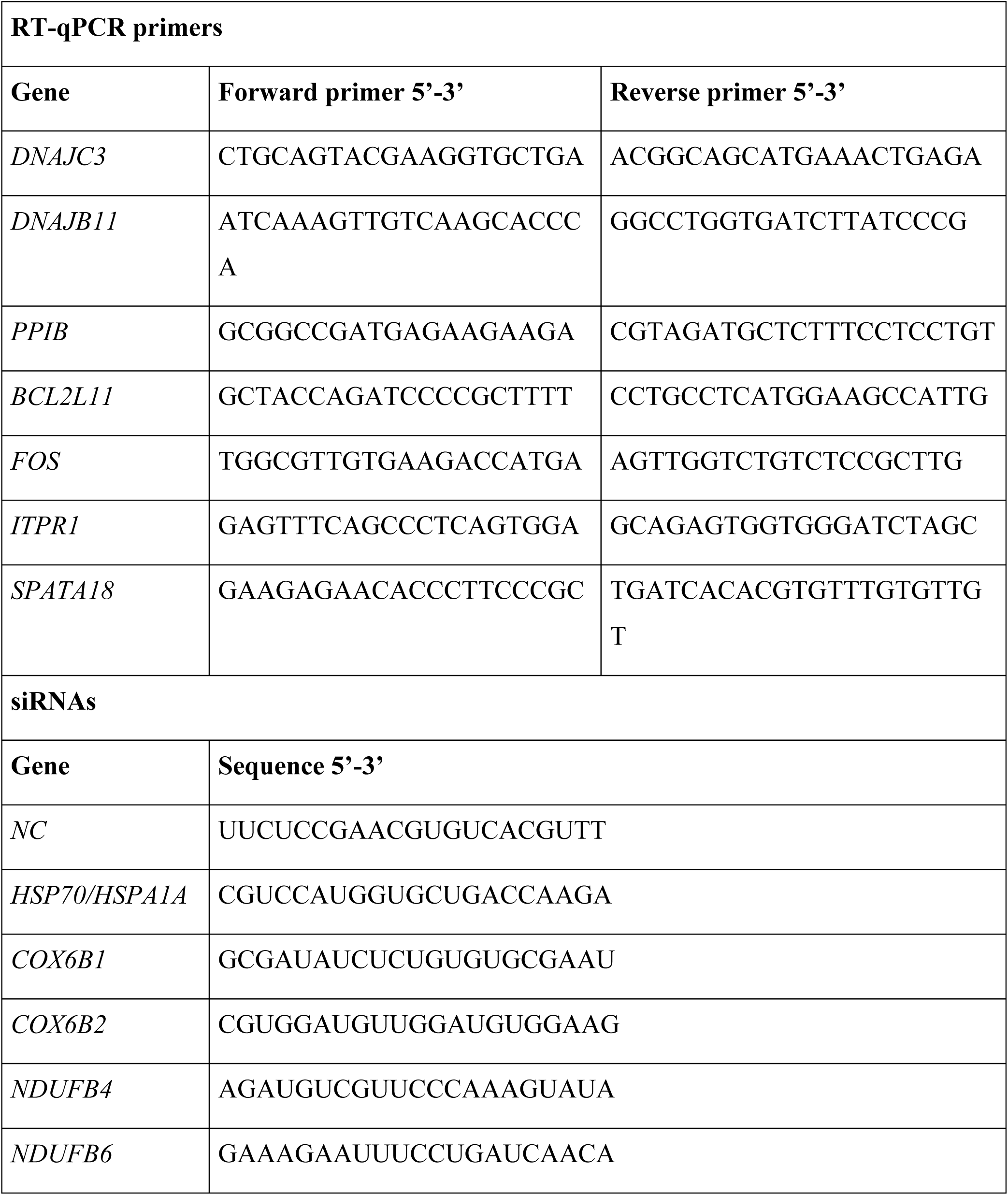
Primer and oligo sequences.

